# Membranes arrest the coarsening of mitochondrial condensates

**DOI:** 10.1101/2025.06.06.658068

**Authors:** Sanjaya Aththawala Gedara, Surya Teja Penna, Marina Feric

## Abstract

Mitochondria contain double membranes that enclose their contents. Within their interior, the mitochondrial genome and its RNA products are condensed into ∼100 nm sized (ribo)nucleoprotein complexes. How these endogenous condensates maintain their roughly uniform size and spatial distributions within membranous mitochondria remains unclear. Here, we engineered an optogenetic tool (mt-optoIDR) that allowed for controlled formation of synthetic condensates upon light activation in live mitochondria. Using live cell super-resolution microscopy, we visualized the nucleation of small, yet elongated condensates (mt-opto-condensates), which recapitulated the morphologies of endogenous mitochondrial condensates. We decoupled the contribution of the double membranes from the environment within the matrix by overexpressing the dominant negative mutant of a membrane fusion protein (Drp1^K38A^). The resulting bulbous mitochondria had significantly more dynamic condensates that coarsened into a single, prominent droplet. These observations inform how mitochondrial membranes can limit the growth and dynamics of the condensates they enclose, without the need of additional regulatory mechanisms.

## Main

Mitochondria are double-membrane bound organelles that form dynamic, tubular networks spanning the cytoplasm. Their membranes help contain multiple copies of their own genome (mtDNA), which encodes key subunits needed for oxidative phosphorylation. The mitochondrial genome is not freely diffuse within the matrix, but is packaged into discrete ∼100 nm nucleoprotein complexes called mt-nucleoids^1^. Under normal conditions, these structures tend to be distributed uniformly throughout the mitochondrial network^2^, yet frequently assemble into large clusters in response to various forms of stress^3,4^. Indeed, there are other examples of distinct nucleic acid and/or protein complexes within the mitochondrial matrix, including mtRNA granules^5–7^, protein translation compartments^8,9^, and mitochondrial stress bodies^8^. However, the mechanisms by which biomolecules organize into such higher-order structures within mitochondria are still largely unclear.

Weak, multivalent interactions among biomolecules can promote the entry into a phase transition, resulting in the formation of condensed liquid-like phases within cells^10^. Often these interactions involve intrinsically disordered regions (IDRs)^11–16^, which can be sufficient to drive phase separation. Indeed, the molecular grammar driving condensation has been well described in the cytoplasm and nucleus^17,18^. However, only recently has phase separation been observed for mt-nucleoids and RNA granules within mitochondria; importantly, the molecular driving forces are largely unexplored^3,19,20^. Moreover, mitochondria represent a unique physicochemical environment in the cell, in part due to their highly membranous structure^21–23^ and high metabolic activities^24–26^, and it remains largely unclear how the mitochondrial physiochemical environment affects the ability of its contents to undergo phase separation.

One tool used to non-invasively study phase separation in live cells involves optogenetics^27–31^. Optogenetic constructs have been engineered by typically fusing an optogenetic protein that dimerizes or oligomerizes in response to light with a sequence capable of undergoing phase separation, such as that of an IDR^27,30–32^. Light activation thus allows spatiotemporal control of condensation, facilitating detailed mapping of cellular phase behavior within a given subcellular environment. Here, we report the development of an optogenetic construct, mt-optoIDR, to directly study phase separation in live mitochondria. With this optogenetic tool targeted to the mitochondria, we report how mitochondrial membrane architecture influences fundamental aspects of phase separation: droplet morphology, dynamics, nucleation and dissolution.

## Results

### mt-optoIDR undergoes a concentration-dependent phase transition in live mitochondria

We adapted an optogenetic approach to induce the formation of condensates in live mitochondria. Our design of an optogenetic construct (mt-optoIDR, Fig. 1a) was based on the established optoIDR constructs used in the nucleus and cytoplasm. Specifically, we retained the optogenetic protein CRY2olig that undergoes reversible protein-protein clustering in response to blue light^31,33^ and its fusion to the red fluorescent marker protein mCherry. Given the unique physicochemical environment of the mitochondrial matrix, we prioritized using an intrinsically disordered region from the mitochondrial proteome that would capture endogenous sequence features. After bioinformatic screening of the mitochondrial proteome (see Methods, Extended Data Fig. 1), we identified the disordered N-terminal region of DDX28 (1 to 127 amino acids) that also contained a mitochondrial targeting sequence (MTS), ensuring our construct would be localized exclusively to the mitochondria. We confirmed the disordered nature using Alphafold2 (Extended Data Fig. 1)^34^. After transient co-expression with a marker to visualize the mitochondrial network (Halo-MTS), mt-optoIDR localized diffusely within the mitochondrial matrix (Fig. 1c, Extended Data Fig. 2a).

**Fig. 1:**
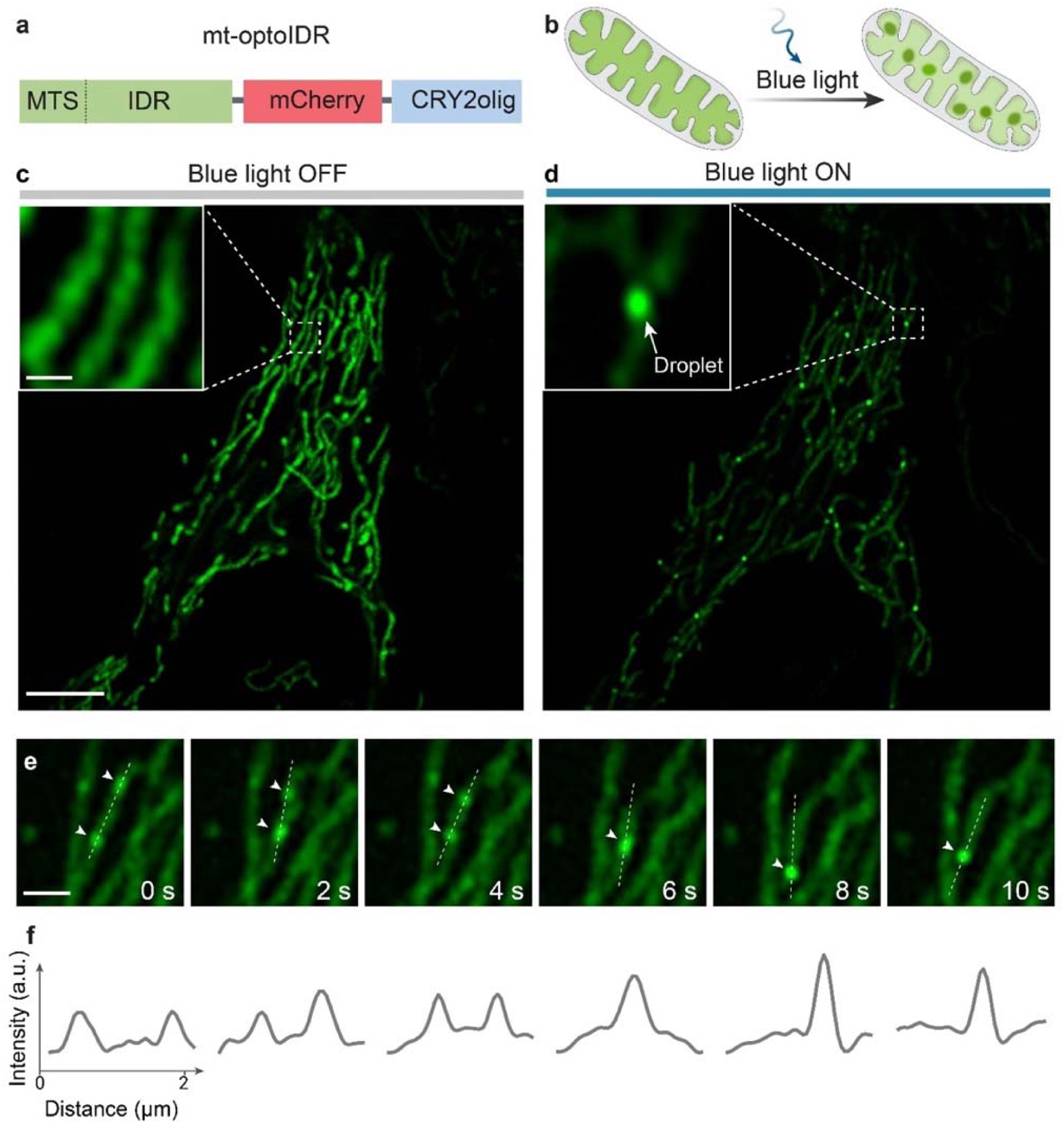
Light-induced phase separation of mt-optoIDR within the mitochondrial matrix of live cells **a**, Schematic of the mt-optoIDR construct containing a mitochondrial targeting sequence (MTS, 1-23) and an intrinsically disordered region (IDR) from DDX28 (24-127 aa) fused to a fluorophore mCherry and an optogenetic protein CRY2olig. **b**, Schematic diagram illustrating light activation experiment using 488 nm laser to induce phase separation of mt-optoIDR within the mitochondrial matrix (see Methods). **c**, Pre-activation (blue light OFF, no droplets) and **d**, post-activation (blue light ON, droplets formed) of HeLa cells expressing mt-optoIDR (green). Puncta in **d** are phase-separated mt-opto-condensates. Inset shows a zoomed in version of the mt-opto-condensate, indicated by the arrow. Scale bars = 5 µm (main) and 0.5 µm (inset). **e**, A liquid-like fusion event of mt-opto-condensate droplets the cell in **c**. Arrowheads point to droplets undergoing coalescence. Scale bar = 1 µm. **f**, Corresponding intensity profiles of the fusion event in **e**. Intensities are normalized based on the first frame. 19 cells were obtained with similar results.

Upon five minutes of blue light activation, mt-optoIDR appeared to undergo phase separation, forming dozens of droplet-like structures (mt-opto-condensates) throughout the mitochondrial network of a HeLa cell (Fig. 1b, d, Extended Data Fig. 2b-d, Supplementary Video 1).

We next performed a series of experiments to confirm that mt-optoIDR exhibited features representative of a classic phase transition. First, we observed that neighboring mt-opto-condensates rapidly fused together, consistent with liquid-like behavior (Fig. 1e-f). Second, the mt-opto-condensates also represented *de novo* phases, as indicated by the lack of partitioning into existing endogenous condensates (Extended Data Fig. 4). Third, we compared droplet formation as a function of expression level or concentration of our construct. We found that for low concentrations mt-optoIDR remained soluble, but above a concentration threshold, formed a second, condensed phase (Extended Data Fig. 3c-e). As a control, mt-opto-condensate formation occurred in the absence of our marker of the mitochondria (Halo-MTS) (Extended Data Fig. 3a-b), but was sensitive to high levels of the mitochondrial marker (Extended Data Fig. 3c). Together, these results show that our optogenetic construct can be used to induce a concentration-dependent phase transition into a condensed liquid-like phase within live mitochondria.

### mt-opto-condensates exhibit narrow size distributions and are elongated in the direction of tubular mitochondria

One prominent feature of the mt-opto-condensates was that they appeared relatively small (Fig. 1, Extended Data Fig. 2, Supplementary Video 1), particularly when compared to reports of opto-droplets formed in the nucleus and cytoplasm^27,31^. We performed quantitative image analysis to measure their size (see Methods) and observed several key features. First, the size distributions of the major and minor droplet axes were non-Gaussian and relatively narrow, ranging from a few hundred nanometers (∼100 nm), suggesting these droplets were limited in their ability to grow into larger droplets. Second, we observed a significant difference between the major and minor axes (p=1.03 × 10^-148^): we estimated the droplet major axis diameter to be 270 ± 50 nm (mean ± s.d, n=533) and droplet minor axis diameter to be 190 ± 20 nm (mean ± s.d, n=533) (Fig. 2a), yielding an aspect ratio (A.R.) of 1.5 ± 0.3 (mean ± s.d, n=533) (Fig. 2b), confirming these condensates were not spherical (A.R. = 1.0) as would be expected for a droplet at equilibrium. Remarkably, these morphologies are roughly consistent with that of endogenous mitochondrial condensates^20,35,36^, indicating the structure of mitochondria may be influencing the morphology of their condensates.

**Fig. 2:**
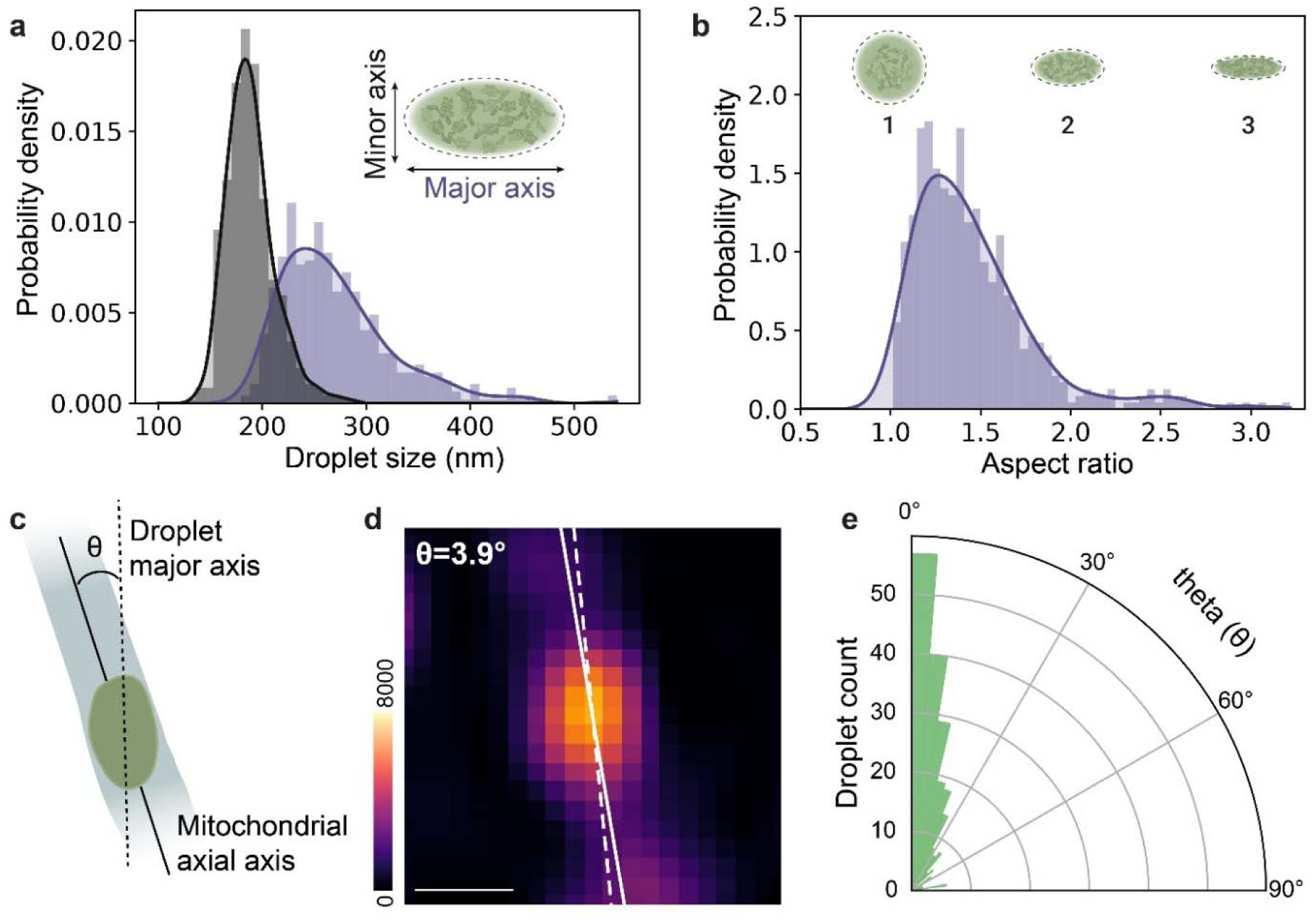
mt-opto-condensates are elongated in the direction of tubular mitochondria. **a**, Quantitative analysis of major (purple) and minor (grey) axes of mt-opto-condensates formed in HeLa cells (n=533 droplets from 14 cells, p-value 1.03 × 10^-148^, Mann-Whitney U test, two sided). Inset includes annotation of major and minor axes. **b**, Aspect ratio changes of mt-opto-condensates (n=533 droplets from 14 cells). Inset includes schematic diagrams illustrating droplets with aspect ratios (AR = 1, 2, and 3) for reference. **c**, Schematic diagram illustrating the angle (θ) between droplet major axis (dotted line) and mitochondrial axial axis (solid line). **d**, An image of the mt-opto-condensate indicating the measured angle, θ. Scale bar = 0.25 µm. **e**, Polar histogram showing the distribution of the measured angle (θ) (n=230 droplets from 14 cells).

We thus reasoned that tubular nature imposed by mitochondrial membranes may play a role in limiting the growth and relaxation of the droplets. If so, the membranes may compress or deform the droplets, resulting in the observed deviation from a spherical shape. To test this hypothesis, we defined a parameter (theta, θ), which is the angle between mitochondrial axial axis and droplet major axis (Fig. 2c-d). If the droplets were being deformed by the membrane, we would expect the droplets to be aligned with the mt-membrane, with correspondingly small theta values. Indeed, the distribution of theta measured for elongated droplets resulted in small angles (θ = 18°±1°, circular mean ± circular s.e.m, n=230), reflecting that the droplets are elongated in the direction of the mitochondrial axial axis of wild-type, tubular mitochondria (Fig. 2e). These results point to the architecture of the mitochondria influencing the morphology of mitochondrial condensates.

To specifically determine the role of the mitochondrial membrane in shaping condensate morphology, we overexpressed the dominant negative mutant of the mitochondrial fission protein Drp1. Drp1^K38A^ is known to lead to hyperfused mitochondria and even more extreme phenotypes, including bulbous structures^37^. Consistent with previous reports^20,37^, the overexpression of Drp1^K38A^ mutant led to the formation of swollen, bulbous mitochondria, allowing us to decouple the contribution of the double membranes from the matrix environment (Extended Data Fig. 5a-c). We noted that there were differences in expression level of our mt-optoIDR construct in tubular and bulbous mitochondria. The bulbous mitochondria showed signs of a stress phenotype, leading to higher levels of the mt-optoIDR construct, roughly twice as concentrated, (p = 0.002, Extended Data Fig. 5d). We repeated the light activation experiments and found a single prominent opto-condensate within each bulbous mitochondrion, indeed reaching considerably larger sizes up to ∼500 nm (Extended Data Fig. 5e). Together, these experiments indicate that mitochondrial membranes can influence the size of condensates.

### Mitochondrial membranes limit the diffusion of mt-opto-condensates

To explain why condensates were forming larger structures in the bulbous mitochondria, we performed time-lapse experiments to capture their dynamics. In wild-type, tubular mitochondria, we noticed that mt-opto-condensates primarily moved significant distances along the direction of the mitochondrial axial axis (dimension *X*, Extended Data Fig. 6a), while being confined to the diameter of the mitochondria (dimension *Y*) (Fig. 3a-c, Supplementary Video. 2,3). We saw significantly more dynamic behavior, irrespective of directionality, in the bulbous mitochondria (Fig. 3d-f, Supplementary Video. 4).

**Fig. 3:**
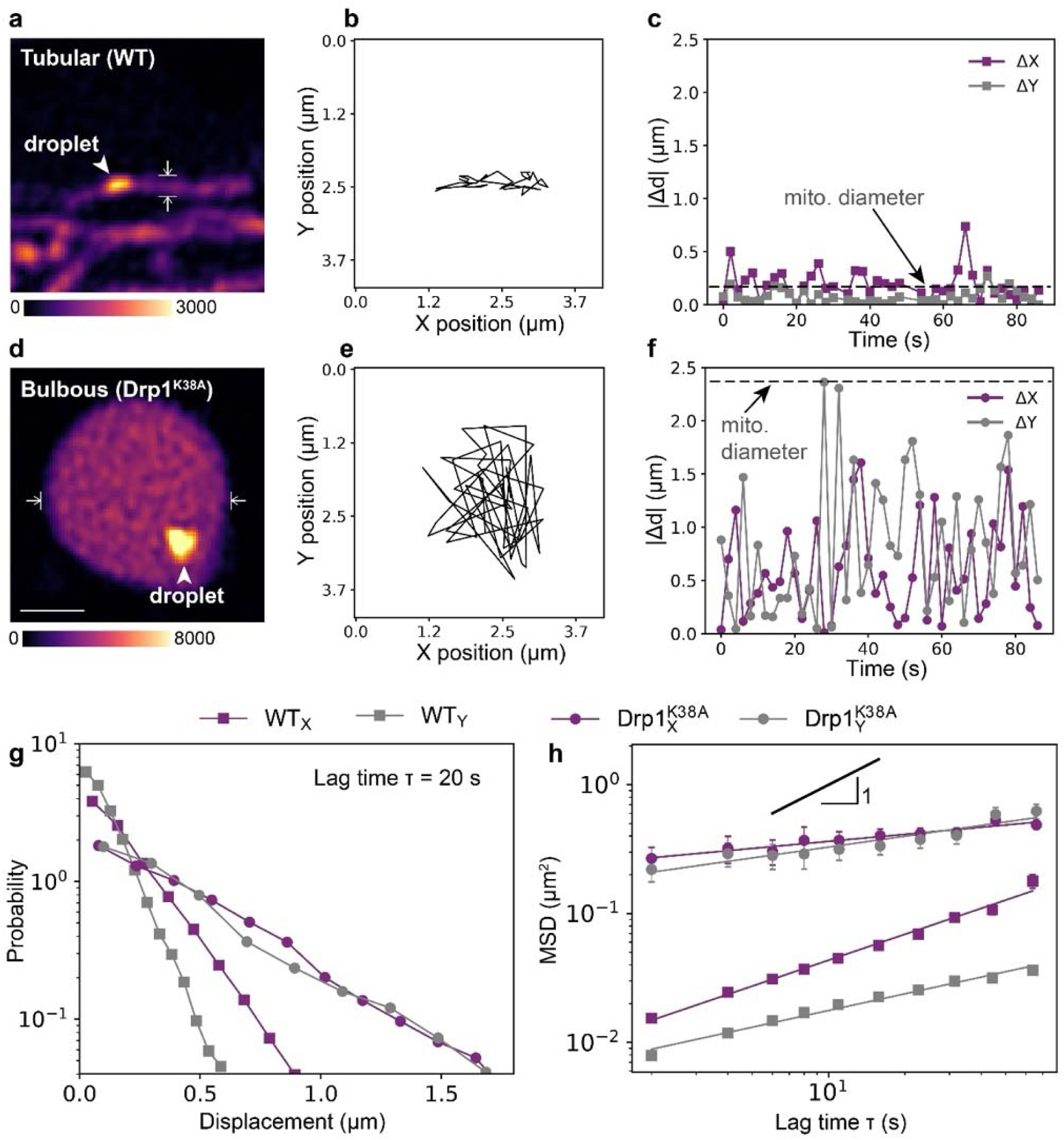
Diffusive motion of mitochondrial condensates is influenced by mitochondrial membrane structure **a,b**, 2D tracking of the mt-opto-condensate in tubular, wildtype mitochondria (image (**a**), track (**b**)). Arrowhead shows the mt-opto-condensate. Flat headed arrows indicate the mitochondrial diameter. **c**, Absolute displacement (Δd) for a lag time, T, of 2 seconds, where Δ*X* (purple) and Δ*Y* (gray) correspond to the mitochondrial axial axis and transverse axis, respectively. **d,e**, 2D tracking of mt-opto-condensate in a bulbous mitochondrion (Drp1^K38A^) (image (**d**), track (**e**)). Arrowhead shows tracked mt-opto-condensate. Flat headed arrows indicate the mitochondrial diameter. **f**, Absolute displacement (Δd) for a lag time, T, of 2 s. **g**, Displacement probabilities of mt-opto-condensates movements in wild-type tubular (square) and bulbous Drp1^K38A^ mitochondria (circles). 2D displacement is decoupled into *X* (purple) and *Y* (gray) dimensions after individual tracks were aligned to *X* axis (see Methods). **h**, Mean Squared Displacement (MSD) of mt-opto-condensates in tubular wild-type and bulbous Drp1^K38A^ mitochondria. Diffusive exponents (α) are a = 0.67±0.06, a = 0.43±0.05, a = 0.18±0.04, and a = 0.28±0.08. The errors are presented as 95% confidence intervals of the standard error of the fit, where n=1,310 and n=25 tracks were identified from 11 wild-type and 10 Drp1^K38A^ transfected HeLa cells, respectively.

To further quantify the constrained diffusion behavior of mt-opto-condensates in wild-type mitochondria, we developed a particle tracking image analysis workflow (see Methods). We extracted the mt-opto-condensate tracks within a tubular mitochondrion and rotated individual tracks to be parallel to the *X* axis in a 2D Cartesian coordinate (Extended Data Fig. 6a-d), such that the new coordinate system had the mitochondrial axial axis as the *X* axis, and the transverse axis as the *Y*-axis (Extended Data Fig. 6a). For example, when absolute displacements (|Δd|) of a mt-opto-condensate were calculated between consecutive frames in a tubular wild-type mitochondrion (Fig. 3a-c), mt-opto-condensates exhibited higher displacements (ΔX >100 nm) along the mitochondrial axial axis compared to the transverse axis (ΔY <100 nm) (Fig. 3c). In comparison, when we performed the same analysis for a mt-opto-condensate in bulbous mitochondria with disrupted membranes, displacements were significantly higher in both dimensions (ΔX, ΔY >>100 nm) yet bound by the diameter of the bulbous mitochondria (Fig. 3d-f). Moreover, the displacements in both *X* and *Y* dimensions were indistinguishable, indicating the mt-opto-condensates were able to freely sample the entire matrix within a bulbous mitochondrion (Fig. 3d-f, Supplementary Video 4). These experiments demonstrate that the motion of mt-opto-condensates is dependent on membrane architecture.

We performed further analysis on the mt-opto-condensate tracks within tubular and bulbous mitochondria. We calculated the displacement probability distributions in both *X* and *Y* dimensions for a given lag time (T = 20 s) (Fig. 3). For Brownian motion in a simple liquid, the displacements in *X* and *Y* dimensions are independent of each other, exhibiting Gaussian distributions. Whereas, for tubular mitochondria, the displacements of mt-opto-condensates in *X* and *Y* were significantly different, where motion in *Y* was largely confined. However, displacements in bulbous mitochondria had broader distributions than for tubular mitochondria, and the distributions were roughly identical in both *X* and *Y* dimensions (Fig. 3g, Extended Data Fig. 6e).

Lastly, we computed the mean squared displacement (MSD) for both wild-type mitochondria and bulbous mitochondria in each dimension (*X*, *Y*). We found power-law behavior across the lag times we experimentally sampled. For both wild-type and bulbous mitochondria, diffusion was highly sub-diffusive as characterized by the diffusive alpha exponent (α). Specifically, MSD for wild-type tubular mitochondria in *X* dimension (along the mitochondrial axial axis) shows anomalous sub-diffusive behavior (α = 0.67±0.06, slope and 95% confidence interval), whereas the diffusive exponent was smaller in the *Y* dimension (transverse axis, α = 0.43±0.05, slope and 95% confidence interval). However, we observed significantly different diffusive behavior in bulbous mitochondria. The MSD values were an order of magnitude larger in value in bulbous mitochondria than in tubular mitochondria, reflecting that the mt-opto-condensates were much more mobile when the membranes were spaced apart. However, bulbous mitochondria had lower diffusive alpha exponents (α), roughly identical for each dimension (α in *X* = 0.18±0.04, α in *Y* = 0.28±0.08, slope and 95% confidence interval), which is characteristic of confined motion. Together, these results show that the diffusion of mt-opto-condensates is ultimately constrained by the imposed membrane-defined boundaries.

### Nucleation, coarsening, and dissolution of mt-opto-condensates

From our global light activation experiments on the entire cell, we observed that the bulbous mitochondria tended to contain a single, prominent mt-opto-condensate, whereas tubular mitochondria were filled with multiple, smaller condensates. We next sought to test if the membrane architecture contributed to the number of droplets formed. We performed a series of local light activation experiments in which we applied light locally to a small region of the cell and imaged concurrently with activation as to capture the nucleation process in real time (see Methods).

Upon local activation in wild-type tubular mitochondria, mt-opto-condensates began to nucleate after ∼100 s of activation and continued to increase in number (Fig. 4a, Extended Data Fig. 7a,c, Supplementary Video 5,6). Similar to the results from our five-minute global activation experiments (Fig. 1, Fig. 2), we did not observe droplets growing beyond ∼100 nm in size, even with longer periods of activation, up to ∼15 minutes. To quantify the nucleation process, we calculated the droplet density (ρ) as a function of time^38^. Upon initiation of liquid-liquid phase separation in classic systems, the droplet density rapidly increases as nucleation occurs followed by a subsequent decrease in cluster number as coarsening processes occur, such as coalescence or Ostwald ripening^31,38^. However, in wild-type tubular mitochondria, the droplet density after ∼15 minutes of activation saturated around 0.15 droplets µm^-2^, likely due to the confined nature of mt-opto-condensates in the mitochondrial matrix. We measured the average nucleation rate (*J*, see Equation 1 in Methods) by taking the slope at t =0 to be 0.0005 droplets µm^-2^ s^-1^, which was defined as the number of visible droplets per unit mitochondrial area per unit time. Moreover, when blue light was turned off, mt-opto-condensates dissolved within several minutes (T = 940 seconds) (Fig. 4b, e, Extended Data Fig. 7b, d, Supplementary Video 7), consistent with a reversible phase transition.

**Fig. 4:**
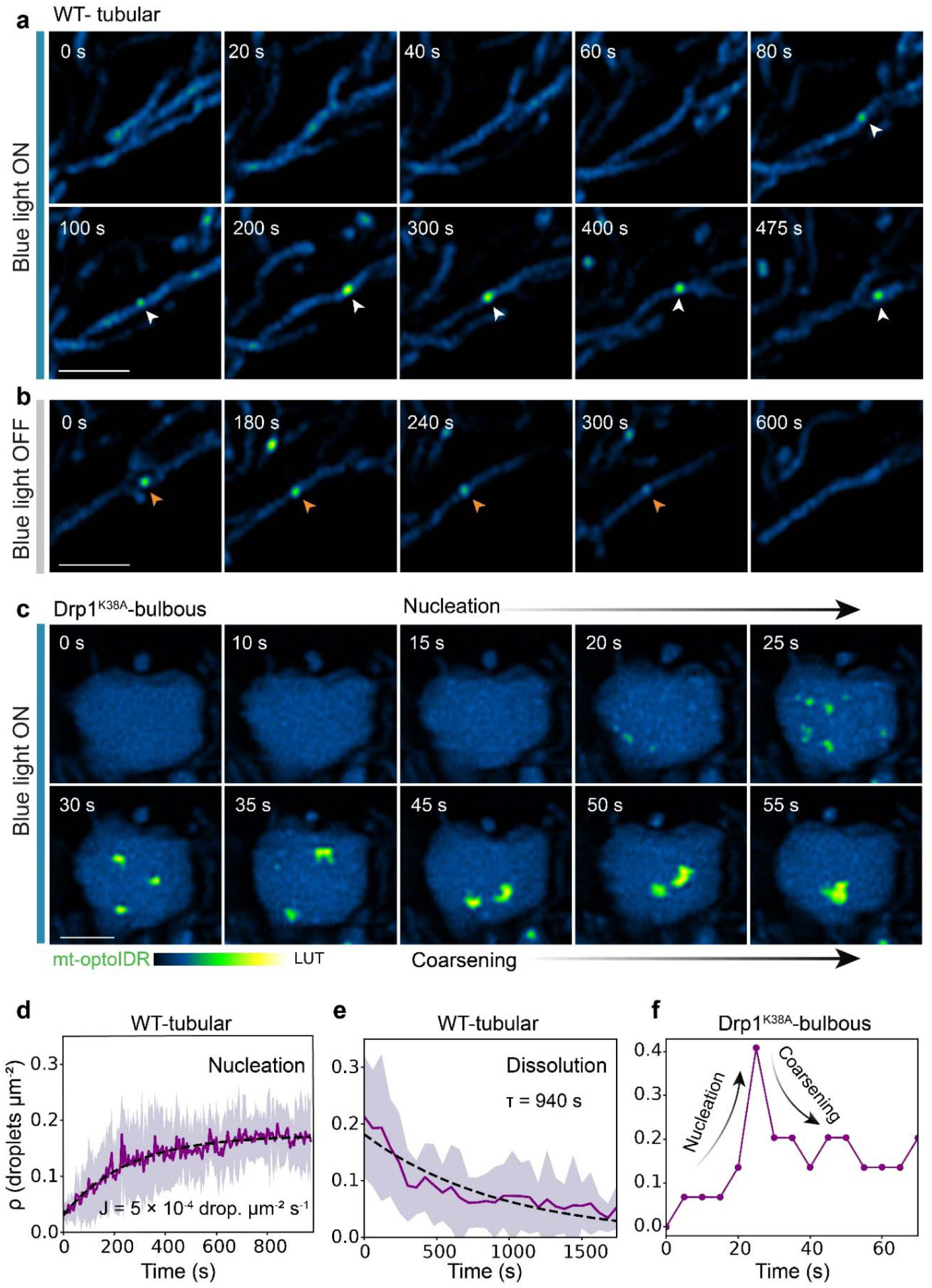
Nucleation and dissolution of mt-opto-condensates. **a**, Local activation (blue light ON) and observation of nucleation of a single mt-opto-condensate within a HeLa cell (see Methods). Arrowheads point to the nucleating droplet. **b**, Removal of blue light and observation of dissolution of a single mt-opto-condensate (blue light OFF). The arrowheads point to the dissolving droplet (same cell as in **a**). Scale bar = 2 µm in **a** and **b**. **c**, Local activation and observation of nucleation and coarsening of mt-opto-droplets in a bulbous mitochondrion (Drp1^K38A^). The first row shows the nucleation process while the second row shows the coarsening of the nucleated droplets. Scale bar = 2 µm. **d,** Droplet density (ρ) as a function of time for condensate nucleation in wild type mitochondria (n=6 cells). The shaded area shows the standard deviation. The dashed line is the fit to equation 2 (see Methods). **e**, ρ as a function of time for condensate dissolution in wild type mitochondria (n=7 cells). The shaded area shows the standard deviation. The dashed line is the fit to equation 3 (see Methods). **f,** Quantification of ρ as a function of time for the bulbous mitochondrion in **c**.

We observed markedly different nucleation dynamics within bulbous mitochondria. We found that after 20 s of activation, numerous small nucleation clusters formed, which rapidly coarsened by fusing together within a particularly large bulbous mitochondrion, leading to a reduction in droplet density (Fig. 4c, Supplementary Video 8). mt-opto-condensates also dissolved within bulbous mitochondria, albeit with slower kinetics than in tubular mitochondria (Extended Data Fig. 8). Together, these experiments demonstrate that mitochondrial membranes act as physical barriers that limit the growth and coarsening of the condensates they contain.

## Discussion

We developed an optogenetic construct, mt-optoIDR to induce phase transitions in live mitochondria. Using mt-optoIDR, we formed *de novo* condensates (mt-opto-condensates) that captured the salient morphological features and dynamic behavior characteristic of endogenous mitochondrial condensates. The dominant negative mutant Drp1^K38A^ allowed us to isolate the contribution of the mitochondrial membrane architecture. Indeed, we found that mitochondrial membranes restricted the diffusion of condensates, preventing their ability to coarsen into prominent droplets.

Endogenous mitochondrial (ribo)nucleoprotein complexes – mt-nucleoids and mtRNA granules – persist as stable droplet-like structures that are known to disassemble *only* if their core nucleic acids are degraded^19,39,40^. One advantage of our optogenetically inducible system is that it allows us to capture the condensation and dissolution processes of condensates within mitochondria live. Phase separation occurs through two mechanism: nucleation and growth where small clusters form that increase in size gradually and spindle decomposition during which large, wide-spread fluctuations in concentration occur before a stable, condensed phase forms^27,38^. Indeed, we observed that our mt-optoIDR nucleates into small clusters that grow into ∼100 nm sized, liquid-like droplets, consistent with nucleation and growth kinetics. We did not observe signs of spinodal-decomposition in live mitochondria, potentially due to the limited expression levels we were able to sample with mt-optoIDR. Nonetheless, upon light-induced nucleation, the mt-opto-condensates were also capable of dissolving when light activation was removed, confirming that biomolecules can undergo reversible phase transitions within live, wild-type mitochondria.

Endogenous mitochondrial condensates are typically small, elongated (A.R. ∼1.5) structures, reaching only a few hundred nanometers^2,20,35,36,41^. Strikingly, our mt-opto-condensates closely mirrored the reported size range with correspondingly elongated aspect ratios, suggesting that the morphology of endogenous mitochondrial condensates need not be actively regulated by the cell but can arise spontaneously due to the biophysical nature of mitochondria. Interestingly, these morphologies differ from the condensates formed in the nucleus and cytoplasm, which can form micron-sized, spherical droplets^27,31^, prompting that the mitochondrial environment may be contributing to the morphology of its condensates. Indeed, we find that mt-opto-condensates are aligned in the direction of tubular, wild type mitochondria, supporting that the three-dimensional structure of mitochondria is the main determinant of condensate morphology. The overall organization is thus likely defined by the interplay between the droplet material properties (e.g. surface tension) and mechanics of the membrane (e.g. bending energies)^42,43^.

Importantly, mitochondrial condensates tend to also be evenly spatially distributed within the mitochondrial network. Indeed, there are multiple mechanisms reported for explaining how mt-nucleoids achieve their uniform spatial distributions, putatively arising from active processes during mitochondrial fission at ER contact sites^44–46^, spontaneous pearling behavior^47^ and/or cristae structure^48–51^. Consistently, the growth of assembled mt-opto-condensates plateaus around ∼100 nm sized droplets, and the resulting structures have significantly limited diffusivities in wild-type mitochondria. These results confirm that condensates become confined within tubular, membranous mitochondria.

Mitochondria can also lose their canonical tubular structure, taking on more bulbous-like morphologies, particularly under pathological conditions. Such bulbous mitochondria have been associated with large clusters of mt-nucleoids and mtRNA granules^3,4,20^. Here, we show that the clustering phenotype of our mt-optoIDR construct arises due to coarsening of condensates in membrane-disrupted mitochondria. Together, our results provide direct evidence that the mitochondrial membranes act as physical barriers restricting the free diffusion of mt-condensates, and in this way, prevent their coarsening. Recent work shows that condensates are known to interact with membranes in other cellular contexts^42,43,52–56^. For example, ER membranes influence the assembly and disassembly of associating condensates^52^, membrane-condensate wetting regulates autophagosome formation^57^, and membranes nucleate signaling components to form productive signaling condensates^58,59^. Our findings show that membranes can also influence droplet morphology by sterically preventing their diffusion and growth. Overall, we find that the kinetics of phase transitions within mitochondria are limited by the architecture of their membranes.

## Statistics and reproducibility

Each experiment had at least three technical replicates (n ≥ 3 cells) with each having multiple mt-opto-condensates, unless otherwise stated. The exact sample sizes in each experiment are reported in the figure legends. Each experiment in the main figures was repeated independently three times on different days. For expression levels and droplet size comparison, the non-parametric, two-tailed Mann-Whitney U test was used since the data do not follow a Gaussian distribution. All the p-values were calculated from the Mann-Whitney U test. p-values are indicated in figure legends.

## Data availability

Source data used to generate figures are provided with this paper through GitHub and FigShare. All data supporting the findings of this study are available from the corresponding author upon reasonable request.

## Code availability

All the analysis scripts used in this study are available through the GitHub webpage at https://github.com/fericlab/mt-opto-IDR.

## Acknowledgments

We thank all the members of Feric laboratory and PSU Center for Eukaryotic Gene Regulation (CEGR) for their feedback and discussion. Research reported in this publication was supported by the National Institute of General Medical Sciences of the National Institutes of Health under award number R35 GM154931 (MF).

## Contributions

M.F. and S.A.G. designed the study, discussed the results and wrote the paper. S.A.G. performed experiments and analyzed data. S.T.P. and S.A.G. wrote code. All authors reviewed the manuscript.

## Supplementary Information

Methods References

Extended Data Figures 1-8 and Figure Legends Supplementary Videos 1-8 and Video Legends Supplementary Tables 1-3

