## Supplementary Information for "Membranes arrest the coarsening of mitochondrial condensates"

**This PDF file includes:**

Methods
References
Extended Data Tables 1 to 3
Extended Data Figures 1 to 8
Supplementary Video Legends

**Other Supplementary Materials for this manuscript include:**

Supplementary Videos 1 to 8

### Methods

#### Bioinformatic analysis

To find an intrinsically disordered region (IDR) endogenous to the mitochondrial proteome, we screened the MitoCarta 3.0 database[1]. We searched sequences from proteins that are known to localize to endogenous mt-condensates (mt-nucleoids or mtRNA granules) that contain an IDR region on the N-terminus immediately succeeding a mitochondrial targeting sequence (MTS). To identify IDR sequence propensity, $D^2P^2$ disorder predictions[2] and AlphaFold structures[3] were used. The presence of the mitochondrial targeting sequence was validated with predictions from Mitofate[4] and TargetP 2.0[5]. After the initial screening as shown in Extended Data Fig 1, four sequences were identified and were cloned (see below). We found best expression of the construct containing the MTS (1-23) and IDR region (24-127) of the mitochondrial dead box helicase 28 (DDX28, 1 to 127 amino acids), which we used in our mt-optoIDR construct. We confirmed the disordered nature of the identified IDR region (24-127) using ColabFold (v1.5.5)[6]. Inputs that deviated from default values include: num_relax = 5, model_type = alphafold2, and relax_max_iterations = 0. Protein structure was viewed using Mol* Viewer[7].

#### Construct design and cloning

Mitochondrial targeting sequence (MTS) (1-23 aa) and intrinsically disordered region (IDR) (24-127 aa) of DDX28 were isolated from total RNA (wild-type primary human fibroblast, HGFDFN168, Progeria Research Foundation[8]) through reverse transcription first-strand cDNA synthesis. The manufacturer's protocol from Thermo Scientific RevertAid Reverse Transcriptase (EP0441) was followed using Oligo(dT)$_{18}$ as primer for the first strand cDNA synthesis, RNase inhibitor (Invitrogen), and dNTP mixture (TaKaRa). From the first strand cDNA library, gene-specific primers were used to perform PCR to amplify the MTS-DDX28(IDR) fragment (forward primer – 5'-GAAACATGGCTCTAACGCGG-3', reverse primer – 5'-GTGCTAGACTGCACGGTTGT-3') using the Primer-BLAST web server[9]. CRY2olig (Addgene plasmid 60032)[10] was cloned into C-terminus of the mCherry in pmCherry-N1 (*TaKaRa,* 632523) (mCherry-CRY2olig). Then MTS-DDX28(IDR) was cloned to the N-terminus of plasmid mCherry-CRY2olig. A linker was placed in between each fragment of MTS-DDX28(IDR), mCherry, and CRY2olig (see Extended Table 1). Fragment assembly was performed using NEBuilder HiFi DNA Assembly kit (see Extended Table 2).

#### Mammalian cell culture

HeLa cells (ATCC, CCL-2, Lot #70046455) were cultured in Dulbecco's Modified Eagle Medium (Gibco, 11960069) at 37°C supplemented with 10% (v/v) Fetal Bovine Serum (FBS) (Gibco, A5256701), and 1% Penicillin /Streptomycin/L-Glutamine (Gibco, 10378-016).

**Transient transfection**

Cells were first seeded in an 8-well imaging chamber (Lab-Tek 155409) on Day 0. On Day 1, cells were transiently transfected at 70-90% confluency with Lipofectamine 3000 transfection kit following manufacturer's protocol (Invitrogen, L3000-001). Briefly, the Lipofectamine-mix had 10 µL Opti-MEM medium (Gibco, REF 31985-062) and 0.3 µL Lipofectamine 3000 (Invitrogen, 100022049). DNA-mix: 0.2 µg DNA (total amount), 10 µL Opti-MEM and 0.4 µL P3000 reagent (Invitrogen, 100022056). 10 µL of DNA-mix and 10 µL of lipofectamine-mix (1:1 ratio) were mixed and incubated for 10 min at room temperature. Then, 20 µL of the DNA-lipid complex (mixed solution) was added to each well of an 8-well imaging chamber. All volumes are presented as per well (0.7 $cm^2$). Volumes were scaled up according to the number of wells transfected during each imaging session. Cells were imaged after two days of transfection (Day 4). All cells were transfected with both mt-optoIDR and Halo-MTS for wild-type cell experiments. Drp1K38A(Addgene plasmid 45161)[11] overexpression was done also in combination with mt-optoIDR and Halo-MTS to induce the bulbous mitochondrial phenotype unless otherwise specified.

**Labeling**

Cells that were expressing Halo-MTS (Addgene plasmid 124315)[12] were labeled with abberior LIVE SiR HaloX (LVSIR-0146-10NMOL, Lot 30707LH-1). Before imaging, cell culture media was removed, and cells were washed with the prewarmed (37°C) live-cell, complete, phenol-red free imaging medium (Gibco 31053036). After that, 200 µL of 0.1 µM staining solution (prepared using 0.1 mM stock solution in DMSO) of HaloX SiR in live-cell imaging media, was added and incubated for 30 min at 37°C, 5% $CO_2$ before imaging. After incubation cells were imaged directly without any washing steps.

**Live cell imaging**

All live cell imaging was performed using a laser scanning confocal microscope equipped with an Airyscan 2.0 detector (Zeiss LSM 980 series). A Plan-Apochromat 63x/1.40 oil DIC M27 oil immersion objective was used to collect images. The microscope is equipped with a built-in incubation chamber maintaining conditions of 5% $CO_2$ and 37°C. Raw images were processed through the Airyscan processing function (Zen Blue software) to achieve super-resolution images, which were used for all subsequent image analysis.

**Global light activation**

Once a single cell was identified**,** global light activation of CRY2olig was first performed on the entire field of view with 488 nm laser (1%) every 5 seconds (total 5 minutes) in confocal mode using the 63x oil objective at 4X zoom with an image size for the activation was 665 × 665 pixels (32.5 µm × 32.5 µm ) and pixel dwell time: 4.1 µs. After the activation, Airyscan super-resolution modes were used to image droplets formed in activated cell, using 561 nm laser (0.5%) and 640 nm laser (0.2%) every 2 seconds for dynamics analysis (total ~6.5 min).

**Local light activation**

Local light activation of CRY2olig was performed with beaching settings in Airyscan line switch mode. 63X oil objective was used at 4X zoom. Bleaching was performed throughout the timelapse, every 5 seconds after each image acquisition with 488 nm laser (1%). Spot size diameter of the region of activation (ROA) = 1 µm circle, scan speed = 1. Image acquisition was done with 561 nm laser (0.5%) and 640 nm laser (0.2%) for a total of ~16.5 min. Light scattering led to diffuse activation surrounding the ROA.

**Immunofluorescence imaging**

Cells were fixed in 4% paraformaldehyde (PFA) (Electron microscopy sciences, 15710) by diluting 16% PFA 1:1 in phosphate-buffered saline (PBS) (MP-Biomedicals,1860454) (8%) and 1:1 (final 4% PFA) directly in cell culture media for 10 min. Then cells were washed with PBS before permeabilization with 0.5% Triton X-100 (Sigma-aldrich, T8787*)* in PBS (v/v) for 10 min at room temperature (RT). Then cells were washed with 0.05% Tween 20 (Sigma-aldrich, P9416*)* in PBS (v/v) three times each two minutes. Blocking was done with 2% Bovine Serum Albumin (BSA, Sigma-aldrich A3294) in 0.05% Tween (w/v) for 30 mins at RT. Blocking agent was removed and specific primary antibodies (diluted in 5% BSA in 0.05% Tween (w/v)) (see Extended Table 3) were incubated for one hour at RT in the dark, followed by three wash steps with 0.05% Tween in PBS, each for two minutes. Then secondary antibodies diluted in the same buffer as primary antibodies were incubated for 1 hour at RT in the dark, followed by three wash steps with 0.05% Tween in PBS for two min each and post fixation in 4% PFA for 10 min. Finally, cells were washed with PBS two times and stored in fresh PBS for imaging.

**Image analysis**

Droplet characterization: For quantitative image analysis, python was used with in-house custom-built scripts. Droplets were located with trackpy (version 0.6.4)[13] package and segmented for further analysis. Characterization of isolated droplets was done by fitting the 2D (*XY*) intensity distribution to a bivariate gaussian. Parameters such as size (Full Width at Half Max, FWHM), and orientation of the droplet were extracted from the fit. All the analysis scripts used are available through the GitHub webpage (https://github.com/fericlab/mt-opto-IDR). For visualization purposes, Fiji (version 2.16.0/1.54p)[14] was used.

Droplet orientation analysis: To determine the orientation of the mitochondrial network, the network was binarized using an adaptive local thresholding. Then the network was skeletonized and isolated the droplet overlapping region of the network to estimate the mitochondrial orientation. A linear fit was performed to the skeletonized mitochondria to estimate the orientation. Then, the orientation of the mitochondria and the orientation of the droplet were used to calculate the angle theta ($\theta$).

MSD analysis: Droplet positions were tracked with trackpy in 2D (*XY*). To isolate independent motion in *X* and *Y*, each droplet track was first aligned to be parallel to the *X*-axis. A linear fit was performed on each track to define the major direction of motion. The angle between *X*-axis and the fit was calculated, allowing us to apply a rotational matrix, which would align the track to the *X*-axis. With the aligned tracks, the Mean Squared Displacement (MSD) was calculated with overlapping lag times, independently for *X* and *Y* dimensions. Only the first 20% of the MSD data was used from each track. To obtain the average MSD, squared displacements were logarithmically binned based on lag times ($\tau$). The error for each data point is reported as standard error of the mean (S.E.M.). The diffusive exponent was obtained from linear fitting of the logarithmic data (log(MSD) versus log($\tau$)). Error of the diffusive exponent is presented as 95% confidence intervals of the slope from the linear fit.

Displacement probabilities: Displacement probabilities were calculated independently for *X* and *Y* dimensions on *X*-aligned tracks.

Nucleation and dissolution: Local activation was used to visualize nucleation of droplets live. For the nucleation and dissolution analysis, 400 pixels by 400 pixel image was isolated centered on the activation region. Droplets were located with trackpy and mitochondrial network area was calculated from the background subtracted image. Nucleation rate, *J,* was calculated at t = 0 s, after fitting to the equation 2[15]:

$$J = \frac{d\rho}{dt} \qquad (1)$$

$$\rho(t) = \rho_0 \left(1 - e^{-\frac{t-t_0}{\tau}}\right) \qquad (2)$$

where $\rho$ is the droplet density, $t$ is time, $\rho_0$ is the initial droplet density, $t_o$ is the time off-set and $\tau$ is the characteristic time. For dissolution, the data was fit to equation 3 to obtain the characteristic time $\tau$:

$$\rho(t) = Ae^{\frac{-t}{\tau}} \qquad (3)$$

where A is a constant.

### Extended Data Tables

Table 1. Linkers used between fragments[16]

| Fragments | Linker (amino acid sequence) |
|---|---|
| mCherry and CRY2olig | GGSGASGGGSGGSGG |
| DDX28(MTS-IDR) and mCherry | GGGGS |

Table 2. Primers used for fragment assembly (UPPERCASE = gene-specific, lowercase = overlap)

| Construct | Fragment | Forward primer (5'-3') | Reverse primer (5'-3') |
|---|---|---|---|
| mCherry-CRY2olig | pmCherryN1fragment | CGGCCGCGACTCTAGATC | CTTGTACAGCTCGTCCATGC |
| mCherry-CRY2olig | CRY2olig fragment | gcatggacgagctgtacaagGGAGGAAGTGGTGCTAGC | atgatctagagtcgcggccgTTATGCTGCTCCGATCATG |
| DDX28(MTS-IDR)-mCherry-CRY2olig | pmCherry-CRY2olig fragment | gcagcggcggcggcggcagcATGGTGAGCAAGGGCGAG | GGTGGCGACCGGTGGATC |
| DDX28(MTS-IDR)-mCherry-CRY2olig | DDX28(MTS-IDR) fragment | gggatccaccggtcgccaccATGGCTCTAACGCGGCCGGTGCGG | GCTGCCGCCGCCGCCGCTGCCCTTAGACGAGAGCTTTCG |

Table 3: Primary and secondary antibodies used in immunofluorescence

| Antibody | Species | 1˚ Ab or 2˚Ab | Dilution | Product information |
|---|---|---|---|---|
| anti-DNA | Mouse | 1˚ Ab | 1:500 | Sigma-Aldrich CBL186 |
| anti-GRSF1 | Rabbit | 1˚ Ab | 1:1000 | Sigma-Aldrich HPA036985 |
| anti-MTCO1 (COX 1) | Mouse | 1˚ Ab | 1:500 | Invitrogen 459600 |
| anti-Tomm20 | Rabbit | 1˚ Ab | 1:500 | Sigma-Aldrich HPA011562 |
| anti-Rabbit-488 | Rabbit | 2˚Ab | 1:1000 | Invitrogen A11008 |
| anti-Mouse-647 | Mose | 2˚Ab | 1:1000 | Invitrogen A32728 |

**Extended Data Figures**
**Extended Data Figure 1**

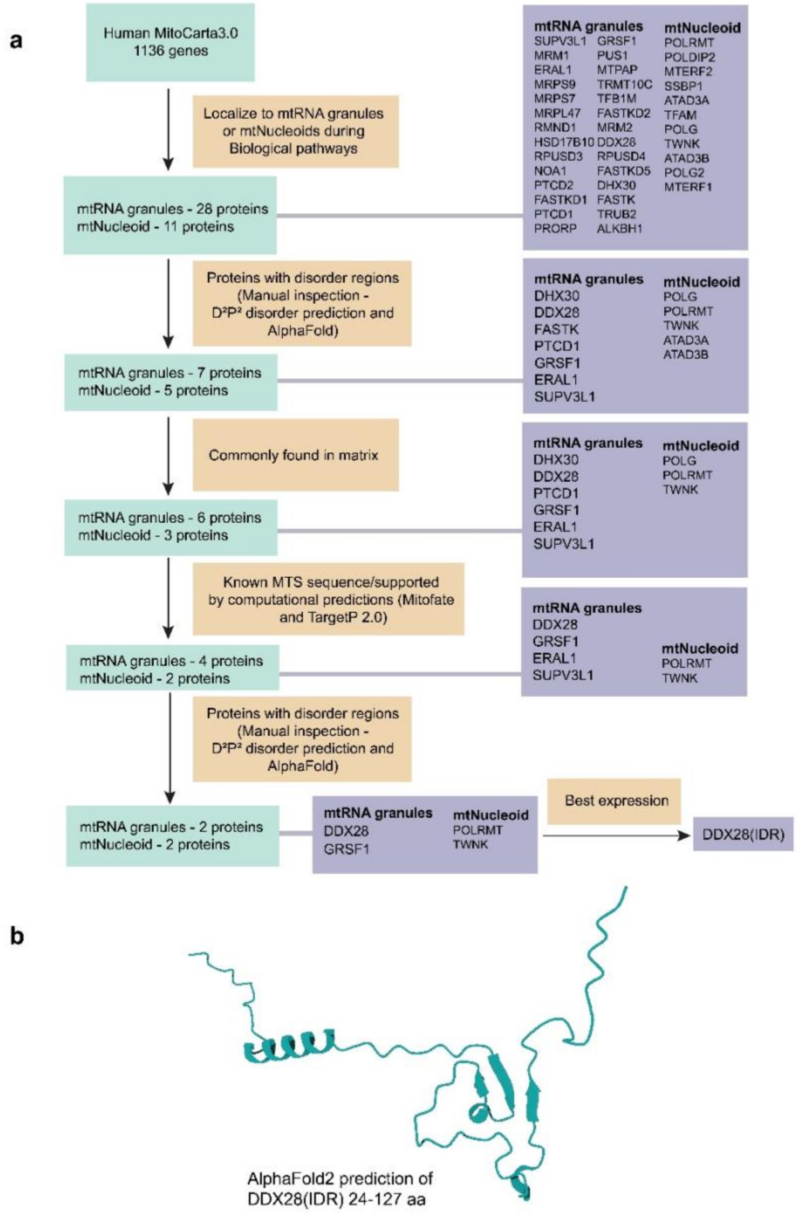


**Extended Data Fig. 1: Bioinformatic analysis pipeline for determining mitochondrial IDR sequence for the mt-optoIDR construct**

**a,** To identify a biologically relevant IDR sequence from the mitochondrial matrix proteome, protein sequences from the MitoCarta 3.0 database were analyzed. Sequences were selected based on disorder predictions from computational models ($D^2P^2$, Alphafold) and presence of an adjacent mitochondrial targeting sequences (MTS). The top four sequences were cloned into mcherry-CRY2olig expression vectors and screened. DDX28(IDR)-mCherry-CRY2olig had the best expression and was used for all subsequent experiments. **b**, Predicted structure of the identified DDX28(IDR) (amino acids 24-127, no MTS) from Alphafold (see Methods).

**Extended Data Figure 2**

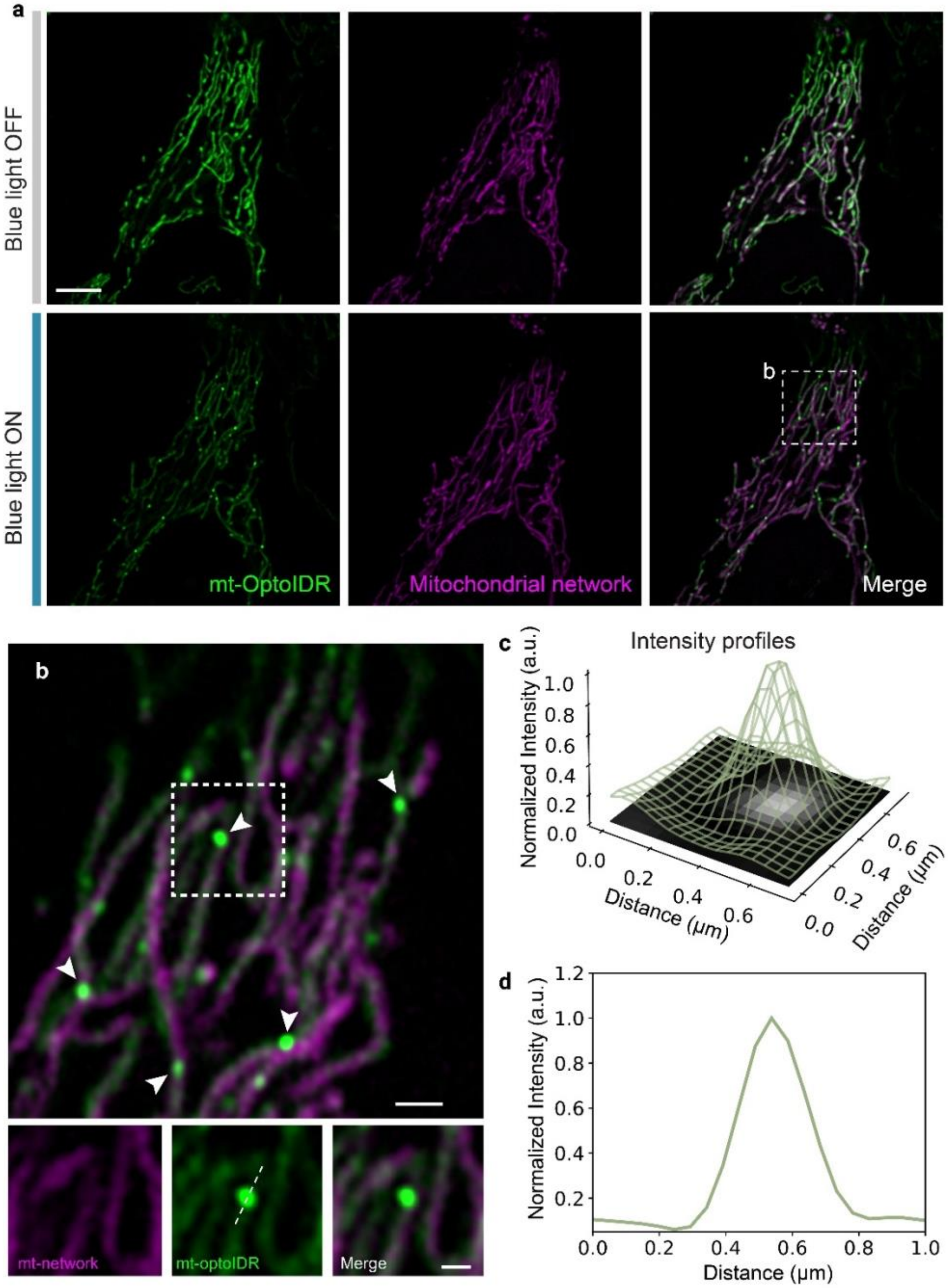


**Extended Data Fig. 2: mt-opto-condensate formation upon light activation**

**a**, A HeLa cell before (top) and after (bottom) light activation. HeLa cells expressing mt-optoIDR construct (green) and mitochondria are visualized using Halo-MTS (magenta). Scale bar = 5 µm. **b**, Zoomed image within the white box in **a** after activation. White arrows point to prominent mt-opto-condensates. Below, zoomed in images of a single droplet. Scale bars = 1 µm (main), 0.5 µm (zoomed). **c**, **d**, Intensity profiles of mt-optoIDR channel of the single droplet in **b** as a 3D representation overlayed on the grayscale image (**c**) and a line profile (**d**). n = 19 cells were imaged with similar results.

**Extended Data Figure 3**

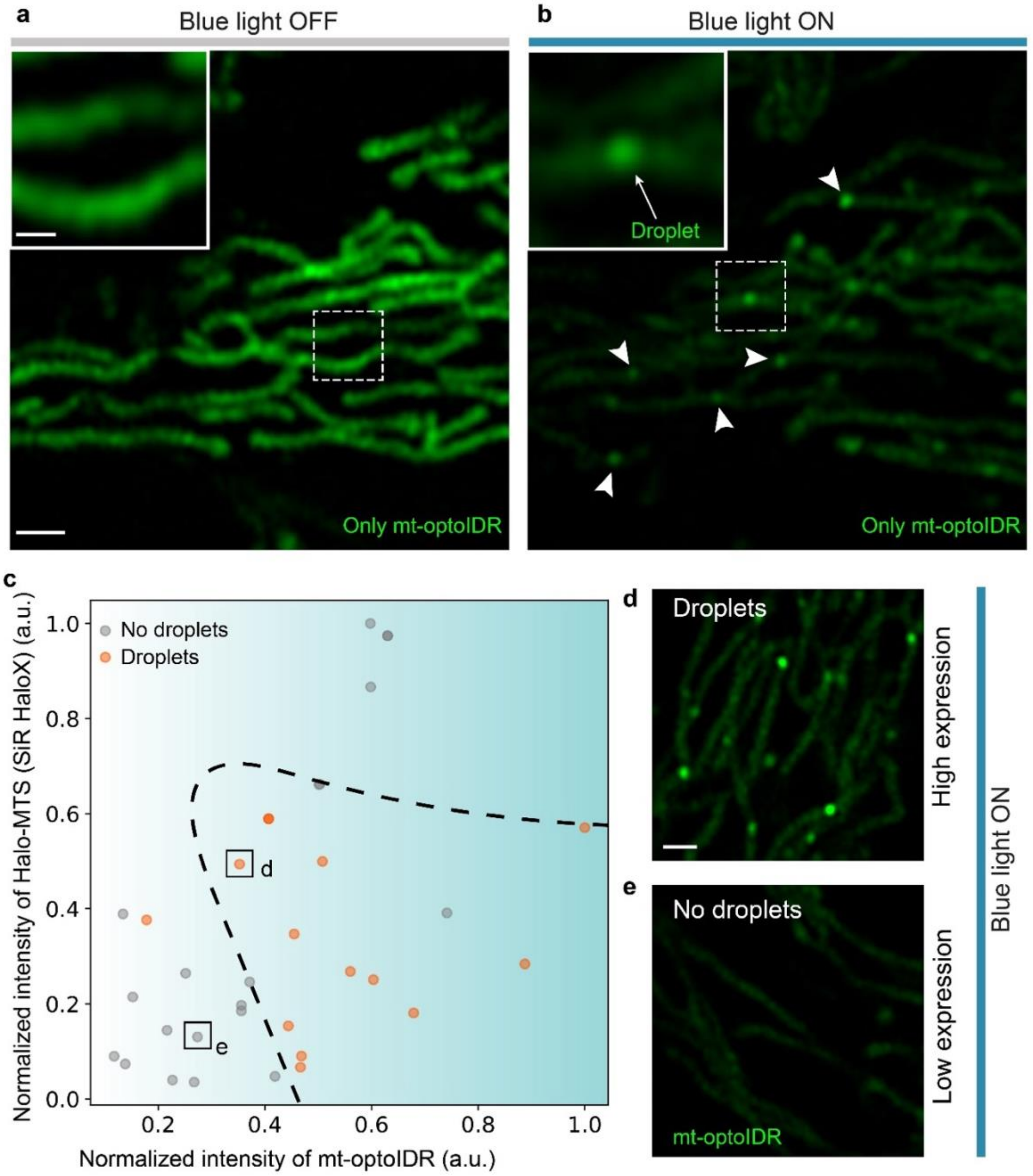


**Extended Data Fig. 3: Concentration dependent phase behavior of mt-optoIDR**

**a, b**, A control cell before (**a**) and after (**b**) light activation when only the mt-optoIDR construct is expressed (i.e. in the absence of Halo-MTS) (n = 3 cells). Arrowheads in **b** point to prominent droplets. Scale bar = 1 µm (main) and 0.5 µm (inset). **c**, Effect of Halo-MTS and mt-optoIDR expression on mt-opto-condensate formation. Different levels of expression are represented as intensities (n = 32 cells). **d, e**, Images visualizing extent of droplet formation upon light activation in cells with high (**d**) or low (**e**) expression of mt-optoIDR. Scale bar = 1 µm.

**Extended Data Figure 4**

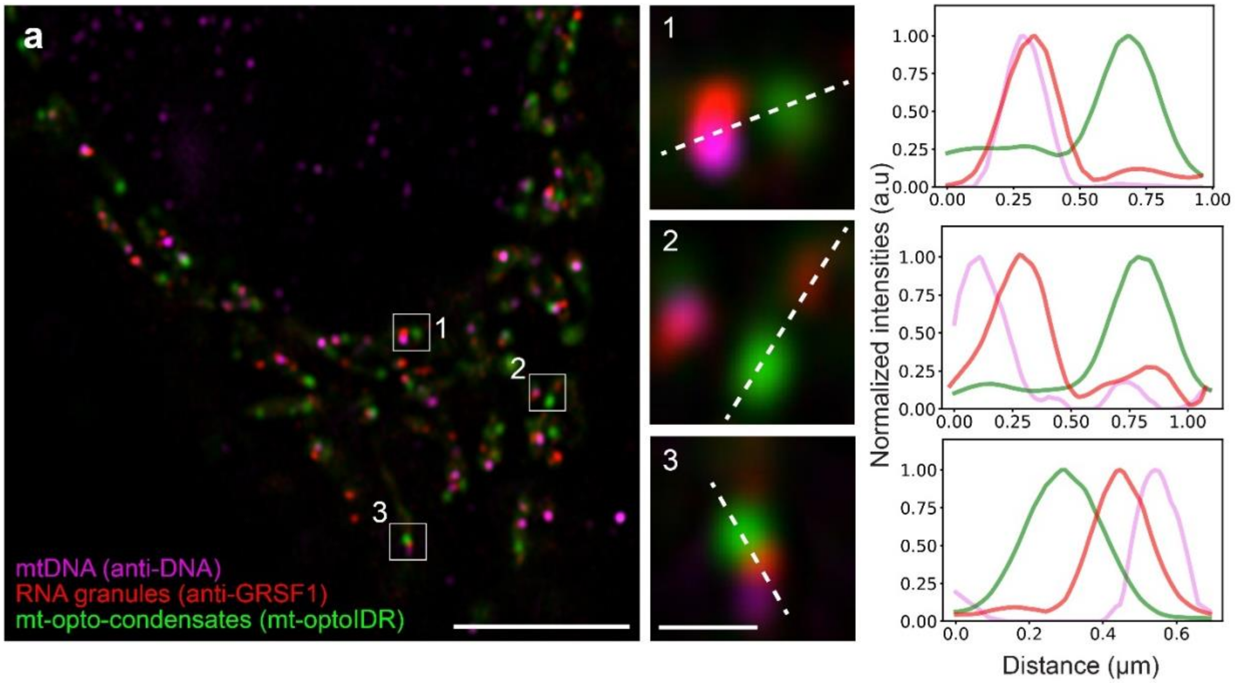


**Extended Data Fig. 4: Spatial distribution of mt-opto-condensates relative to mt-nucleoids and mtRNA granules**

**a**, A representative image showing mt-nucleoids (anti-DNA, magenta), mtRNA granules (anti-GRSF1, red), and mt-opto-condensates (mt-optoIDR, green) (n = 5 cells). The second column shows zoomed images of areas 1, 2, and 3. The third column is the corresponding intensity line profiles for each channel. Scale bar = 5 µm (main) and 0.5 µm (zoomed).

**Extended Data Figure 5**

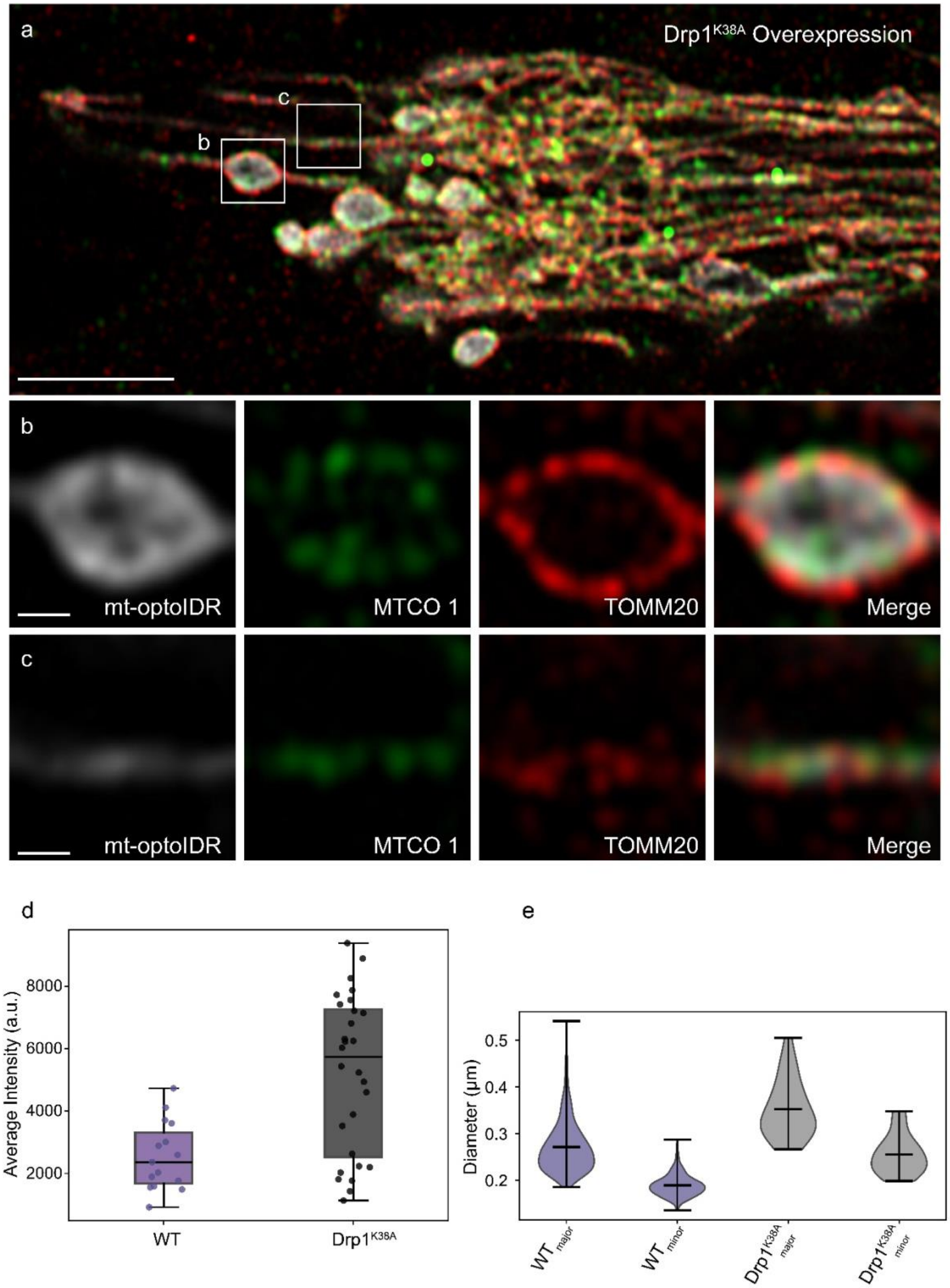

**Extended Data Fig. 5: Drp1$^{K38A}$ overexpression yields large bulbous mitochondria with altered membrane structure**

**a**, A representative immunofluorescence image of a non-activated HeLa cell overexpressing Drp1$^{K38A}$ and labelled mt-optoIDR construct (gray), inner mitochondrial membrane (anti-MTCO1, green), and outer mitochondrial membrane (anti-TOMM20, red) (n = 6 cells). Scale bar = 5 µm. **b-c**, Zoomed in images of bulbous (**b**) and tubular (**c**) mitochondria from main image (**a**). Scale bar = 0.5 µm. **d**, Distribution of expression levels (normalized intensity) of mt-optoIDR in wildtype mitochondria (n=15 cells) versus Drp1K38A mutant bulbous mitochondria (n=28 bulbous mitochondria) (p=0.002, Mann-Whitney U test). **e**, Size comparison (major and minor diameters) of mt-opto-condensates in wild-type tubular mitochondria and Drp1$^{K38A}$ bulbous mitochondria. For Drp1$^{K38A}$ bulbous droplets, the time average is reported for 20 frames in each droplet (WT n=533, Drp1$^{K38A}$ n=24). Statical analysis: p-value = $2.8 \times 10^{-5}$ for $WT_{major}$ vs $Drp1^{K38A}{}_{major}$, p-value = $4.0 \times 10^{-7}$ for $WT_{minor}$ vs $Drp1^{K38A}{}_{minor}$, Mann-Whitney U test, two-sided.

**Extended Data Figure 6**

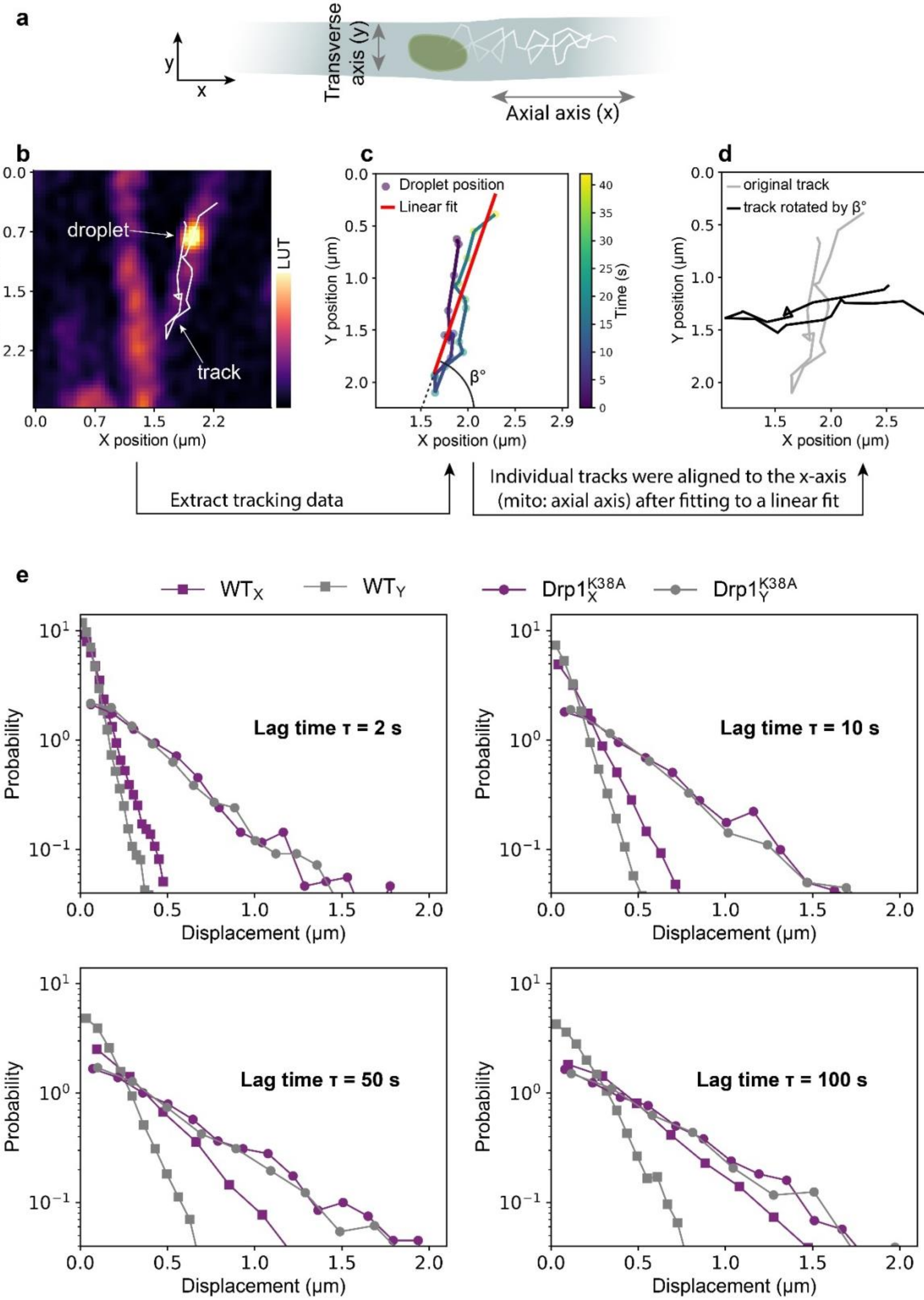

**Extended Data Fig. 6: Analysis of mt-opto-droplets trajectories**
**a**, Schematic diagram of condensate two-dimensional (2D) diffusion within a tubular mitochondrion with annotated mitochondrial axial axis (*X*), and transverse axis (*Y*). **b**, Image of mt-optoIDR condensates in Hela cell with the trajectory overlayed (white track) . **c**, The trajectory from **b** was plotted on an *XY* coordinate system and a linear fit was overlayed. The angle beta (β), was determined relative to the linear fit and the horizontal *X* axis. Heatmap shows the time progression. **d**, Trajectory from **c** (gray) and the rotated trajectory (black). The rotated trajectory was used for all subsequent analysis **e**, Probability distributions for displacement (*X* purple, *Y* gray) at different lag times ($\tau$ = 2, 10, 50, and 100 seconds) in tubular wild type (squares) and bulbous (circles) mitochondria.

**Extended Data Figure 7**

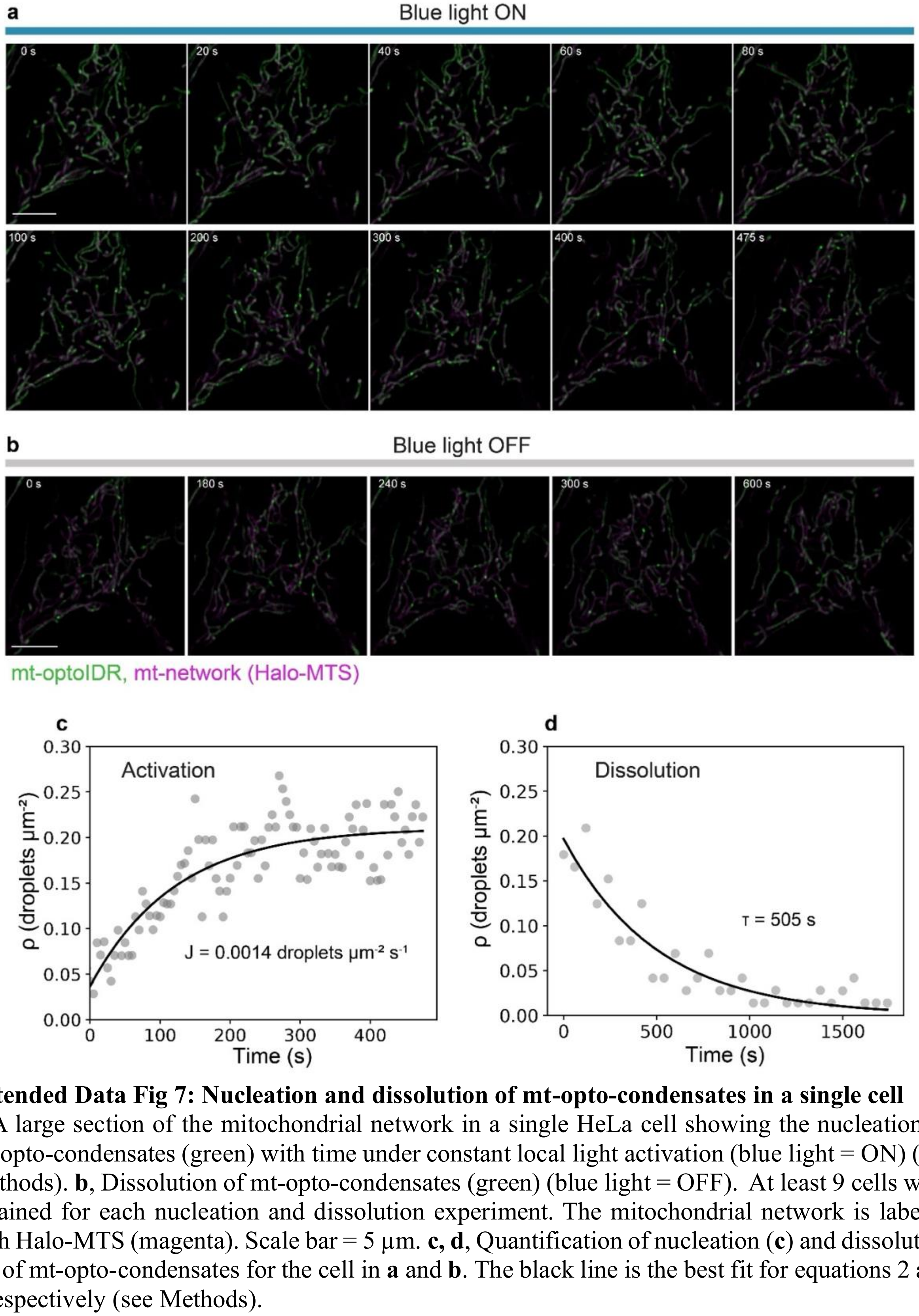


**Extended Data Fig 7: Nucleation and dissolution of mt-opto-condensates in a single cell**
**a**, A large section of the mitochondrial network in a single HeLa cell showing the nucleation of mt-opto-condensates (green) with time under constant local light activation (blue light = ON) (see Methods). **b**, Dissolution of mt-opto-condensates (green) (blue light = OFF). At least 9 cells were obtained for each nucleation and dissolution experiment. The mitochondrial network is labeled with Halo-MTS (magenta). Scale bar = 5 μm. **c, d**, Quantification of nucleation (**c**) and dissolution (**d**) of mt-opto-condensates for the cell in **a** and **b**. The black line is the best fit for equations 2 and 3 respectively (see Methods).

**Extended Data Figure 8**

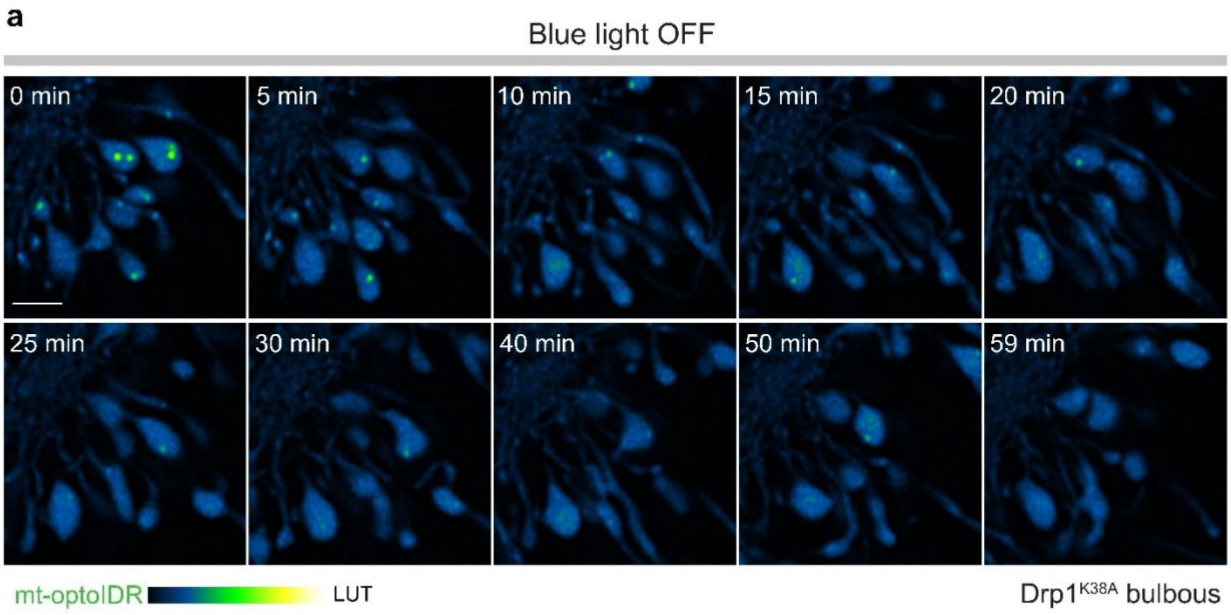


**Extended Data Fig. 8: Dissolution of mt-opto-condensates in bulbous mitochondria**
**a,** A large section of the mitochondrial network in a HeLa cell with Drp1$^{K38A}$ overexpression enriched in bulbous mitochondria showing mt-opto-condensates dissolution (blue light = OFF). At least 9 cells were imaged with the same dissolution behavior. The heatmap shows the mt-optoIDR intensity. Scale bar = 2 µm.

**Supplementary Video Legends**

**Supplementary Video 1: Global light activation results in phase separation of mt-optoIDR construct (green) into liquid-like droplets within mitochondria of live HeLa cells.** The inset shows a zoomed-in version of an individual condensate. Time is reported as seconds. Scale bar = 5 µm.

**Supplementary Video 2: Diffusion of mt-opto-condensates within wild-type, tubular mitochondria.** The white arrowhead points to an individual droplet. Time is reported as seconds. Scale bar = 1 µm. Intensity is represented by the heat-map.

**Supplementary Video 3: Diffusion of mt-opto-condensates (green) within wild-type, tubular mitochondria (magenta).** White arrowheads direct the movements of droplets. The mitochondrial network is labeled with Halo-MTS (magenta). Time is reported as seconds. Scale bar = 1 µm.

**Supplementary Video 4: Diffusion of a mt-opto-condensate within a bulbous mitochondrion**. The bulbous phenotype was induced by $Drp1^{K38A}$ overexpression. Time is reported as seconds. Scale bar = 1 µm. Intensity is represented by the heat-map.

**Supplementary Video 5: Nucleation of mt-optoIDR construct into a droplet within mitochondria upon local light activation**. mt-optoIDR intensity is represented by a heatmap. The mitochondrial network is labeled with Halo-MTS (magenta). The blue square (right top) indicates blue light activation. Time is reported as seconds. Scale bar = 1 µm.

**Supplementary Video 6: Nucleation of mt-optoIDR construct into droplets (green) within mitochondria (magenta) upon local activation.** A large portion of the mt-network of a single HeLa cell is shown. The inset zooms in on the nucleation of a single droplet. The white arrowhead points to the nucleating droplet. The mitochondrial network is labeled with Halo-MTS (magenta). The blue square (right top) indicates blue light activation. Time is reported as seconds. Scale bar = 5 µm.

**Supplementary Video 7: Dissolution of mt-opto-condensates (green) in mitochondria (magenta)**. A large portion of the mt-network of a single HeLa cell is shown. mt-network was imaged without blue light activation, which led to the dissolution of condensates. The inset zooms in on a single droplet dissolving. The white arrowhead points to the droplet. mt-network is labeled with Halo-MTS (magenta). Time is reported as seconds. Scale bar = 5 µm.

**Supplementary Video 8: Nucleation and coarsening of mt-opto-condensates in a large bulbous mitochondrion**. This phenotype was induced by $Drp1^{K38A}$ overexpression. mt-optoIDR intensity is represented as a heatmap. The mitochondrial network is labeled with Halo-MTS (magenta). The blue square (right top) indicates blue light activation. Time is reported as seconds. Scale bar = 1 µm.
